# Efficient use of genomic information for sustainable genetic improvement in small cattle populations

**DOI:** 10.1101/617464

**Authors:** J. Obšteter, J. Jenko, J. M. Hickey, G. Gorjanc

**Affiliations:** Department of Animal Science, Agricultural Institute of Slovenia, Hacquetova ulica 17, 1000 Ljubljana, Slovenia; Geno Breeding and A.I. Association, Storhamargata 44, 2317 Hamar, Norway; The Roslin Institute and Royal (Dick) School of Veterinary Studies, University of Edinburgh, Easter Bush, Midlothian, EH259RG, United Kingdom; Biotechnical Faculty, University of Ljubljana, Jamnikarjeva 101, 1000 Ljubljana, Slovenia

**Keywords:** small population, sustainability, genomic selection, optimum contribution selection

## Abstract

This paper compares genetic gain, genetic variation, and the efficiency of converting variation into gain under different genomic selection scenarios with truncation or optimum contribution selection in a small dairy population by simulation. Breeding programs have to maximize genetic gain but also ensure sustainability by maintaining genetic variation. Numerous studies showed that genomic selection increases genetic gain. Although genomic selection is a well-established method, small populations still struggle with choosing the most sustainable strategy to adopt this type of selection. We developed a simulator of a dairy population and simulated a model after the Slovenian Brown Swiss population with ~10,500 cows. We compared different truncation selection scenarios by varying i) the method of sire selection and their use on cows or bull-dams, and ii) selection intensity and the number of years a sire is in use. Furthermore, we compared different optimum contribution selection scenarios with optimization of sire selection and their usage. We compared the scenarios in terms of genetic gain, selection accuracy, generation interval, genetic and genic variance, the rate of coancestry, effective population size, and the conversion efficiency. The results show that early use of genomically tested sires increased genetic gain compared to progeny testing as expected from changes in selection accuracy and generation interval. A faster turnover of sires from year to year and higher intensity increased the genetic gain even further but increased the loss of genetic variation per year. While maximizing intensity gave the lowest conversion efficiency, a faster turn-over of sires gave an intermediate conversion efficiency. The largest conversion efficiency was achieved with the simultaneous use of genomically and progeny tested sires that were used over several years. Compared to truncation selection optimizing sire selection and their usage increased the conversion efficiency by either achieving comparable genetic gain for a smaller loss of genetic variation or achieving higher genetic gain for a comparable loss of genetic variation. Our results will help breeding organizations to implement sustainable genomic selection.

## INTRODUCTION

This paper compares genetic gain, genetic variation, and the efficiency of converting variation into gain under different genomic selection scenarios in a small dairy cattle population with truncation or optimum contribution selection by simulation. Genomic selection has profoundly changed dairy cattle breeding programs (Schaeffer, 2006; Garcia-Ruiz et al., 2016; Wiggans et al., 2017). It has doubled the rate of genetic gain through decreased generation interval, increased selection accuracy for young animals, increased selection intensity, and identification and management of recessive lethal alleles (Cole, 2015; Garcia-Ruiz et al., 2016; Wiggans et al., 2017). The prerequisite for these gains is a large number of genotyped animals, which is an issue for small populations (Thomasen et al., 2014; Jenko et al., 2017; Ducrocq et al., 2018), though this problem can be addressed with international training populations (Jorjani, 2012; Liu, 2013; Vandenplas et al., 2017). An effective implementation also requires an optimal use of genomic selection for different groups of animals (Thomasen et al., 2014). Further, small populations struggle to maximize selection intensity due to a limited number of animals and limited resources, but also due to genetic drift and related genetic variation issues, which can be enhanced with intense and rapid genomic selection (Falconer and Mackay, 1996; Gorjanc et al., 2018).

Breeding programs aim to maximize genetic gain. Previous studies compared the conventional progeny testing with genomic pre-selection prior to progeny testing or direct genomic selection for widespread use without progeny testing (de Roos et al., 2011; Lillehammer et al., 2011; Pryce et al., 2010). These studies reported up to 30% increase in genetic gain with the genomic pre-selection and up to 195% increase with the direct genomic selection. Thomasen et al. (2014) deterministically evaluated hybrid schemes that use both progeny and young genomically tested sires in populations of different size. They concluded that genomic selection gives higher genetic gain than conventional progeny testing irrespective of population size, but that the hybrid schemes maximize annual monetary genetic gain when a population is small and accuracy of genomic selection is low.

Breeding programs also have to maintain genetic variation to ensure long-term sustainability. This is especially important for small populations, since they have to be competitive in the international market to justify the national breeding program. While short-term success depends on the genetic gain in the next few generations, long-term success depends also on maintenance of sufficient genetic variation to ensure a stable rate of genetic gain (Woolliams et al., 2015). Studies on the effect of genomic selection on genetic variation have had contradictory results. For example, Lillehammer et al. (2011) and Pryce et al. (2010) reported a decreased rate of coancestry per year, while de Roos et al. (2011) reported that it depends on the proportion of genetic variation captured with markers and a breeding program design. Genomic selection has a potential to decrease the rate of coancestry due to a more accurate estimation of Mendelian sampling terms for young animals, which enables differentiation of sibs and avoidance of their co-selection (Daetwyler et al., 2007). Balancing short- and long-term success can be further enhanced with the optimum contribution selection (Woolliams et al., 2015).

Although genomic selection is a well-established method, small populations still struggle with choosing a sustainable strategy. The right strategy should ensure short- and long-term success as well as being economically and logistically viable. To address some of these issues this study evaluates different genomic breeding program designs for a small dairy population with a focus on selection and usage of sires and how this affects changes in genetic gain and genetic variation.

## MATERIAL AND METHODS

We compared conventional and different genomic breeding program designs in a small dairy cattle population with simulation. Altogether we compared twenty-two scenarios. In fifteen scenarios we used truncation selection with five different selection criteria to choose sires for the insemination of cows and bull-dams. Additionally, we tested each of the sire selection criterion scenarios within three sire usage scenarios that varied the number of sires and the period of their usage. To maximize genetic gain for a given loss in genetic variation we compared the truncation selection scenarios with seven optimum contribution selection scenarios where we varied balance between genetic gain and maintenance of genetic variation. We compared all the scenarios in terms of genetic gain, genetic variation, and efficiency of converting genetic variation into gain.

### Simulation

We developed a simulator of a realistic dairy population. The simulator is a Python wrapper around the simulation program AlphaSim (Faux et al., 2016; Hickey and Gorjanc, 2012), the genetic evaluation program blupf90 (Misztal et al., 2002), and the optimum contribution selection program AlphaMate (Gorjanc and Hickey, 2018). The simulator is driven by a set of parameters describing a dairy breeding program, including the percentage of animals selected at each stage and in each selection path, age at selection, selection criterion (pedigree or genomic), the number of progeny per sire, the number of years a group of animals is used, and the number of years a simulation is run. These parameters allow the simulation of relevant dairy breeding programs. In each year the simulator generates phenotypic data, estimates breeding values, culls, selects and mates animals, and generates progeny - including their pedigree and genotypic data.

### Population

The simulated population mimicked the Slovenian Brown Swiss population of ~30,000 animals of which ~10,500 are cows. The simulation started with a coalescent process to generate whole-genome sequence for ten cattle-like chromosomes with 10^8^ base pairs per chromosome, mutation rate of 2.5 x 10^−8^, recombination rate of 1.0 x 10^−8^ and historical effective population size in line with the estimates for dairy cattle (Villa-Angulo et al., 2009; Hickey and Gorjanc, 2012). We randomly sampled segregating sequence variants to construct a set of 10,0000 causal variants (1,000 per chromosome) and two distinct sets of 20,000 marker variants (2,000 per chromosome). We used the two sets of marker variants to create two SNP arrays, one was used for genomic selection and the other for monitoring “neutral” diversity. We sampled the effects of causal variants from a normal distribution with a variance that gave a trait with the heritability of 0.25 in the base population. Subsequently we initiated a base population by randomly allocating simulated genomes to animals, which were further allocated to different categories (male and female calves, cows, bull-dams, young bulls, AI bulls and natural service bulls) to initiate a dairy breeding program. We have then run a conventional breeding program with selection on phenotype based estimated breeding values for 20 years, followed by a further 20 years of different scenarios described below. We repeated simulation of the base population and each scenario 20 times.

We generated 4,320 female calves every year of which we removed a random 3.7% due to stillbirths and early deaths, and a further 7.5% due to other losses, for example, reproductive issues (Figure S1). The remaining heifers were inseminated in the second year and became cows in the third year. In each subsequent lactation we culled 20% of the starting number of cows at random and all remaining cows after the fourth lactation. This scheme totaled to about 10,500 active cows per year. During the first lactation we assigned 43 cows with the highest estimated breeding values as bull-dams and used them for three lactations, which gave us 129 active bull-dams per year (Figure S1). Every year we inseminated the best 90 bull-dams with relevant sires to generate elite male selection candidates.

We selected sires based on genomic or progeny tests. Every year 45 elite male calves were tested following one of three scenarios: a) progeny test with a pre-selection based on pedigree prediction (PT), b) progeny test with a pre-selection based on genomic test (GT-PT) or c) genomic test (GT). With the PT scenario, 8 out of 27 calves were chosen for progeny test based on pedigree prediction in their second year, while the remaining 19 calves were used in natural service (Figure S2). With the GT-PT scenario 8 out of 45 calves were chosen for progeny test based on genomic test. With the PT and GT-PT scenario 5 out of 8 progeny tested bulls were selected as sires based on estimated breeding value in their sixth year (Figure S2). With the GT scenario 5 out of 45 genomically tested calves were directly selected as sires and were used for insemination from their second year onwards. Unselected genomically tested calves in all genomic scenarios were used as natural service sires (Figure S3).

Bull dams were inseminated with selected AI bulls only. For the insemination of cows, AI sires contributed 400 doses of semen per year when 5 sires where used for 5 years and 2,000 doses per year when 5 sires where used for 1 year or 1 sire was used for 5 years; natural service sires contributed 27 doses; and young bulls (where applicable) contributed 250 doses. The expected proportion of offspring of natural service sires therefore ranged between 7 and 17%.

### Breeding value estimation

We estimated breeding values with the pedigree model (Henderson, 1984) or the single-step genomic model (Legarra et al., 2009) using the blupf90 program with default options (Misztal et al., 2002). In genomic breeding scenarios we assumed an initial reference population of about 11,000 cows and 100 progeny tested sires. This mimicked the availability of international genomic evaluation in Brown Swiss (Jorjani, 2012). We updated the reference population each year by replacing the oldest cows with about 2,000 new cows and elite male selection candidates. Variance components were assumed known and set to simulated values.

### Breeding scenarios

We created different truncation selection scenarios by varying i) the method of sire selection and their use on cows or bull-dams, and ii) selection intensity and the number of years a sire is in use. Furthermore, we created different optimum contribution selection scenarios with optimization of sire selection and their usage.

#### Truncation selection

The scenarios that varied the selection of sires in combination with their use on cows or bull-dams were: i) PT scenario used PT sires for the insemination of cows and bull-dams, ii) GT-PT scenario used GT-PT sires for the insemination of cows and bull-dams, iii) GT-C scenario used GT sires for the insemination of cows and GT-PT sires for the insemination of bull-dams, iv) GT-BD scenario used GT sires for the insemination of bull-dams and GT-PT sires for the insemination of cows, and v) GT scenario used GT sires for the insemination of both cows and bull-dams. The GT-C and GT-BD scenarios are also referred to as the hybrid scenarios.

The scenarios that varied selection intensity and the number of years a sire is in use were: i) select five sires every year and keep them in use for five years (5 sires/year, use 5 years), ii) reduce generation interval by using five sires for one year only (5 sires/year, use 1 year) and iii) maximize selection intensity by selecting only one sire and use it for five years (1 sire/year, use 5 years).

#### Optimum contribution selection

We have optimized sire selection and usage with optimum contribution selection (Woolliams et al., 2015) using the AlphaMate program (Gorjanc and Hickey, 2018). Every year we have added the 45 genotyped elite male calves to the pool of sires selected in the previous year with a limit of 5 years for sire usage. We then optimized their contributions while fixing female (heifers’ and cows’) contributions to one progeny per female. After optimization we randomly paired the optimized male contributions with the fixed female contributions. Inputs for optimum contribution selection were estimated breeding values and a coancestry matrix (Woolliams et al., 2015) from the genomic single-step model (Legarra et al., 2009). We optimized contributions with different emphasis on genetic gain versus group coancestry using the target degrees of the angle between the truncation selection solution and an optimum contribution solution (Kinghorn, 2011). For example, target degrees of 0 maximize genetic gain by selecting only one male, while target degrees of 90 solely minimize group coancestry. We evaluated a range of target degrees and reported results for 45, 50, 55, 60, and 75 degrees.

### Analysis

We compared the scenarios in terms of genetic gain, selection accuracy, generation interval, genetic and genic variance, the rate of coancestry, effective population size, and the efficiency of converting genetic variation into genetic gain. Genetic gain was expressed as a deviation from average true breeding values of the individuals in the first year of comparison in the units of genetic standard deviation. Selection accuracy was measured with the Pearson correlation between the true and estimated breeding values. Calibration of estimated breeding values (bias) was measured with the coefficient of regression of true breeding values on estimated breeding values. Generation interval was computed as the average age of the parents at the birth of their selected offspring. Genetic variance measured variance of true breeding values. Genic variance measured variance of true breeding values under the assumption of no linkage between causal loci. The rate of coancestry per year was calculated from pedigree or genomic information. The pedigree coancestry was computed following Wright (1922) from which the rate of coancestry (ΔC_P_) was estimated as one minus the exponent of the coefficient of regression of log(C_P,t_) on year of birth (Pérez-Enciso, 1995). The genomic coancestry was computed based on the direct link with heterozygosity, Het_t_ = Het_o_(1 - C_t_) (Falconer and Mackay, 1996). We computed heterozygosity separately for causal, marker, and neutral loci. We regressed log(C_t_) on the year of birth to estimate the rate of coancestry for causal loci (ΔC_Q_), marker loci (ΔC_M_), and neutral loci (ΔC_N_). Effective population size (N_e_) was estimated for every measure of the rate of coancestry as 1/(2ΔC). Finally, the conversion efficiency was measured with the coefficient of regression of the achieved genetic gain on the loss of genic standard deviation (Gorjanc et al., 2018). This metric quantifies the genetic gain achieved in units of genic standard deviation when all variation is converted into gain or lost due to drift. Results are presented as the mean of 20 replicates for each scenario on a per year or cumulative basis. The progeny testing breeding program with 5 sires selected per year and used for 5 years was the baseline for comparison.

## RESULTS

The results compare different breeding scenarios for a small dairy cattle population in terms of genetic gain, genetic variation, and the efficiency of converting genetic variation into genetic gain. The early use of genomically tested sires increased genetic gain compared to progeny testing. A faster turnover of sires from year to year and higher intensity increased the genetic gain even further but increased the loss of genetic variation per year. The conversion efficiency increased with the simultaneous use of genomically and progeny tested sires. Maximizing intensity resulted in the lowest effective population size and the lowest conversion efficiency. A faster turn-over of sires decreased the conversion efficiency to an intermediate degree. Compared to truncation selection optimizing male contributions increased the conversion efficiency by either achieving comparable genetic gain for a smaller loss of genetic variation or achieving higher genetic gain for a comparable loss of genetic variation.

### Genetic gain

Early use of genomically tested sires, their faster turn-over and higher intensity of selection increased genetic gain. This is shown in Table 1, which presents genetic gain by breeding program and by sire selection and their usage scenario. Genomic pre-selection for progeny testing increased genetic gain by 36% compared to the baseline. Genomic selection of sires for a direct insemination of cows or bull-dams increased genetic gain respectively by 62% or 68%, and by 94% when used for both, cows and bull-dams. Reducing the use of the selected sires from 5 years to 1 year further increased genetic gain, between 10% and 142% compared to the baseline. Reducing the number of selected sires per year from 5 to 1 and using that sire for 5 years also increased genetic gain, between 21% and 124% compared to the baseline, but not compared to the scenario where 5 selected sires per year were used for 1 year. These genetic gains were a direct function of realized generation intervals (Table S1) and selection accuracies (Table S2). Table S2 also reports the calibration (bias) of estimated breeding values.

**Table 1:** Genetic gain in genetic standard deviation units by breeding program and by sire selection and their usage scenario.

| Breeding program | Sire selection and usage |  |  |
| --- | --- | --- | --- |
|  | 5 sires/year, use 5 years | 5 sires/year, use 1 year | 1 sire/year, use 5 years |
| PT | 2.50<sub>0.22</sub><sup>a, A</sup> | 2.75<sub>0.19</sub><sup>a, B</sup> | 3.03<sub>0.11</sub><sup>a, C</sup> |
| GT-PT | 3.41<sub>0.14</sub><sup>b, A</sup> | 3.96<sub>0.17</sub><sup>b, B</sup> | 3.84<sub>0.13</sub><sup>b, B</sup> |
| GT-C | 4.05<sub>0.15</sub><sup>c, A</sup> | 4.65<sub>0.21</sub><sup>c, B</sup> | 4.82<sub>0.21</sub><sup>c, B</sup> |
| GT-BD | 4.20<sub>0.19</sub><sup>c, A</sup> | 4.56<sub>0.25</sub><sup>c, B</sup> | 4.51<sub>0.20</sub><sup>d, B</sup> |
| GT | 4.84<sub>0.26</sub><sup>d, A</sup> | 6.04<sub>0.27</sub><sup>d, C</sup> | 5.60<sub>0.27</sub><sup>e, B</sup> |

PT = conventional progeny testing; GT-PT = genomic pre-selection of bulls for progeny testing, GT-C = genomic selection of sires for the insemination of cows; GT-BD = genomic selection of sires for the insemination of bull-dams; GT = genomic selection of sires for the insemination of cows and bull-dams. Subscript numbers indicate standard deviation across simulation replicates. Lower-case letters denote statistically significant differences between breeding programs and upper-case letters between sire selection and usage scenarios.

### Genetic and genic standard deviation

Early use of genomically tested sires, their faster turn-over and higher intensity of selection decreased genetic variation. This is shown in Figure 1, which presents genic and genetic standard deviation by breeding program and by sire selection and their usage scenario. The genic and genetic standard deviations are expressed as the percentage change to the baseline that had in the final year genic standard deviation of 0.97 and genetic standard deviation of 0.94. Genomic pre-selection for progeny test did not significantly change genic standard deviation. Genomic selection of sires for a direct insemination of cows or bull-dams reduced genic standard deviation between 1.3% and 2.5%. Reducing the number of years sires were used from 5 to 1 further reduced genic standard deviation, between 0.9% and 5.0% compared to the baseline. Increasing selection intensity, by selecting only 1 sire per year instead of 5, reduced genic standard deviation even further, between 3.0 and 10.3%. We observed a similar trend in the reduction of genetic standard deviation as for genic standard deviation, but the reductions were overall larger and had higher variation between simulation replicates.

**Figure 1:**
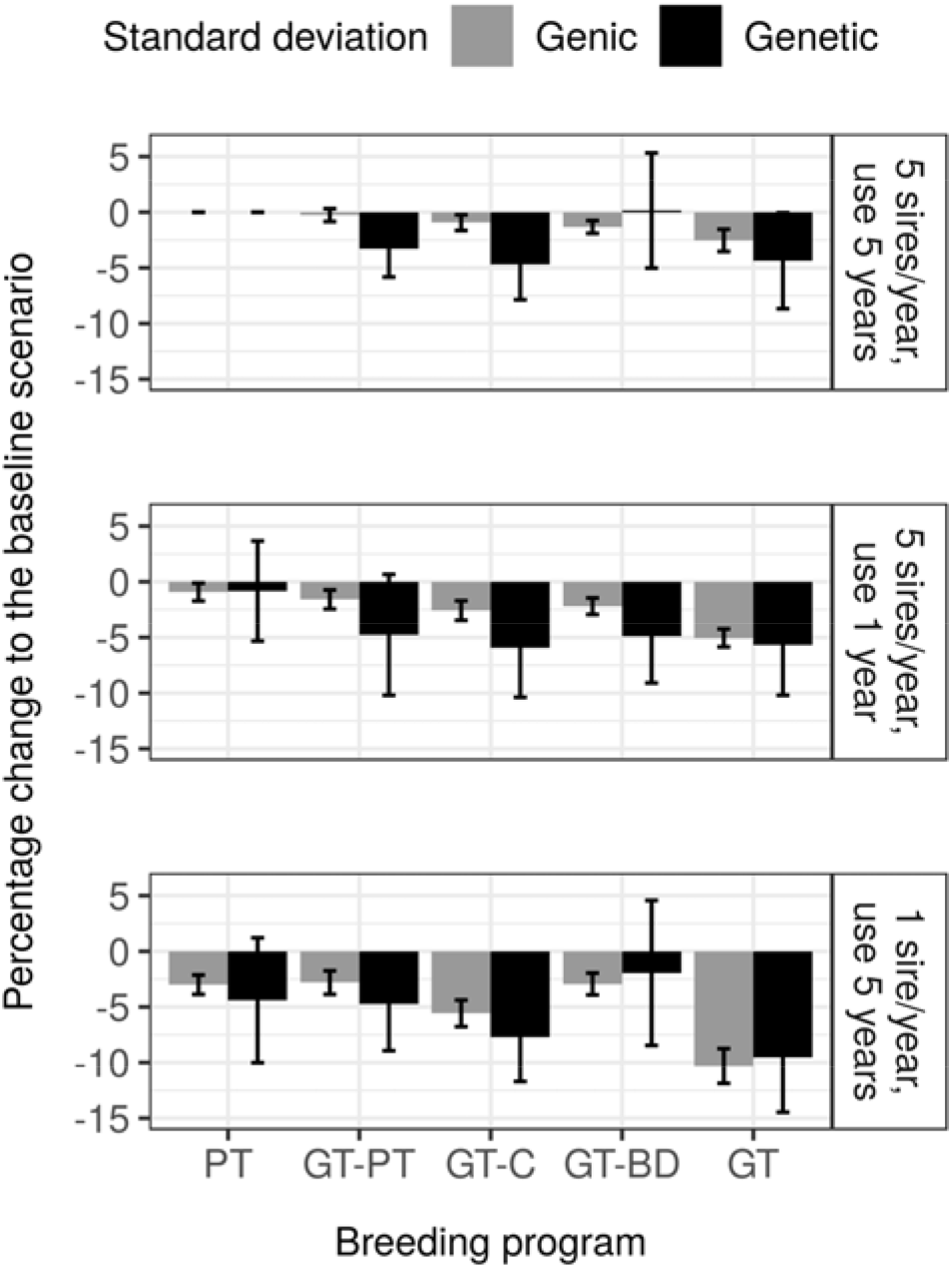
Genic and genetic standard deviation by breeding program and by sire selection and their usage scenario expressed as percentage change to the baseline that had in the final year genic standard deviation of 0.97 and genetic standard deviation of 0.94. PT = conventional progeny testing; GT-PT = genomic pre-selection of bulls for progeny testing, GT-C = genomic selection of sires for the insemination of cows; GT-BD = genomic selection of sires for the insemination of bull-dams; GT = genomic selection of sires for the insemination of cows and bull-dams.

### Effective population size

Early use of genomically tested sires and increased selection intensity decreased effective population size. This is shown in Table 2, which presents effective population size at causal loci by breeding program and by sire selection and their usage scenario. Genomic pre-selection for progeny testing did not significantly change the effective population size. Inseminating cows, bull-dams or both with young genomically tested sires decreased effective population size respectively by 25%, 31%, and 48%. Reducing the years the sires are used from 5 to 1 did not significantly change effective population size for any of the corresponding breeding scenarios. In contrast, reducing the number of sires selected per year from 5 to 1 and using that sire for 5 years decreased effective population size for all scenarios. The decrease ranged from 42% (with genomic pre-selection for progeny testing) to 79% (when both cows and bull-dams were inseminated with one genomically tested sire) compared to the baseline. These results were qualitatively the same as results for the effective population sizes at marker loci used for genomic selection or at “neutral” loci (results not shown).

**Table 2:** Effective population size at causal loci by breeding program and by sire selection and their usage scenario.

| Breeding<br>program | Sire selection and their usage |  |  |
| --- | --- | --- | --- |
|  | 5 sires/year, use 5 years | 5 sires/year, use 1 year | 1 sire/year, use 5 years |
| PT | 172<sub>48</sub><sup>a, A</sup> | 184<sub>57</sub><sup>a, A</sup> | 96<sub>20</sub><sup>a, B</sup> |
| GT-PT | 159<sub>43</sub><sup>a, A</sup> | 146<sub>40</sub><sup>b, A</sup> | 99<sub>20</sub><sup>a, B</sup> |
| GT-C | 129<sub>29</sub><sup>b, A</sup> | 124<sub>32</sub><sup>bc, A</sup> | 64<sub>11</sub><sup>b, B</sup> |
| GT-BD | 119<sub>27</sub><sup>b, A</sup> | 113<sub>24</sub><sup>c, AB</sup> | 93<sub>24</sub><sup>a, B</sup> |
| GT | 90<sub>14</sub><sup>c, A</sup> | 72<sub>10</sub><sup>d, A</sup> | 38<sub>6</sub><sup>b, B</sup> |

PT = conventional progeny testing; GT-PT = genomic pre-selection of bulls for progeny testing, GT-C = genomic selection of sires for the insemination of cows; GT-BD = genomic selection of sires for the insemination of bull-dams; GT = genomic selection of sires for the insemination of cows and bull-dams. Subscript numbers indicate standard deviation across simulation replicates. Lower-case letters denote statistically significant differences between breeding programs and upper-case letters between sire selection and usage scenarios.

### Conversion efficiency

The greatest efficiency of converting genetic variation into gain was achieved with the simultaneous use of genomically and progeny tested sires that were used over several years. This is shown in Table 3, which presents the conversion efficiency by breeding program and by sire selection and their usage scenario. This measure indicates long-term genetic gain in standard deviation units when all genic variance will be exhausted and is calculated by regressing the achieved genetic gain on the lost genic variance over the 20 years of selection, which we graphically represent in Figure 2 to complement the Table 3. Compared to the baseline, the introduction of genomic selection increased the conversion efficiency. The highest increase, 31%, was achieved with the genomic pre-selection for progeny testing. Genomic selection of sires for the insemination of cows or bull-dams increased the conversion efficiency respectively by 28% or 22%. Genomic selection of sires for the insemination of both cows and bull-dams did not significantly increase the conversion efficiency compared to the baseline. Reducing the usage of sires from 5 years to 1 year decreased the conversion efficiency, except for the two scenarios with the highest genetic gain, that is, when using genomically tested sires for the insemination of bull-dams or all females. Reducing the number of selected sires per year to 1 and using it for 5 years reduced conversion efficiency furthermore.

**Table 3:**
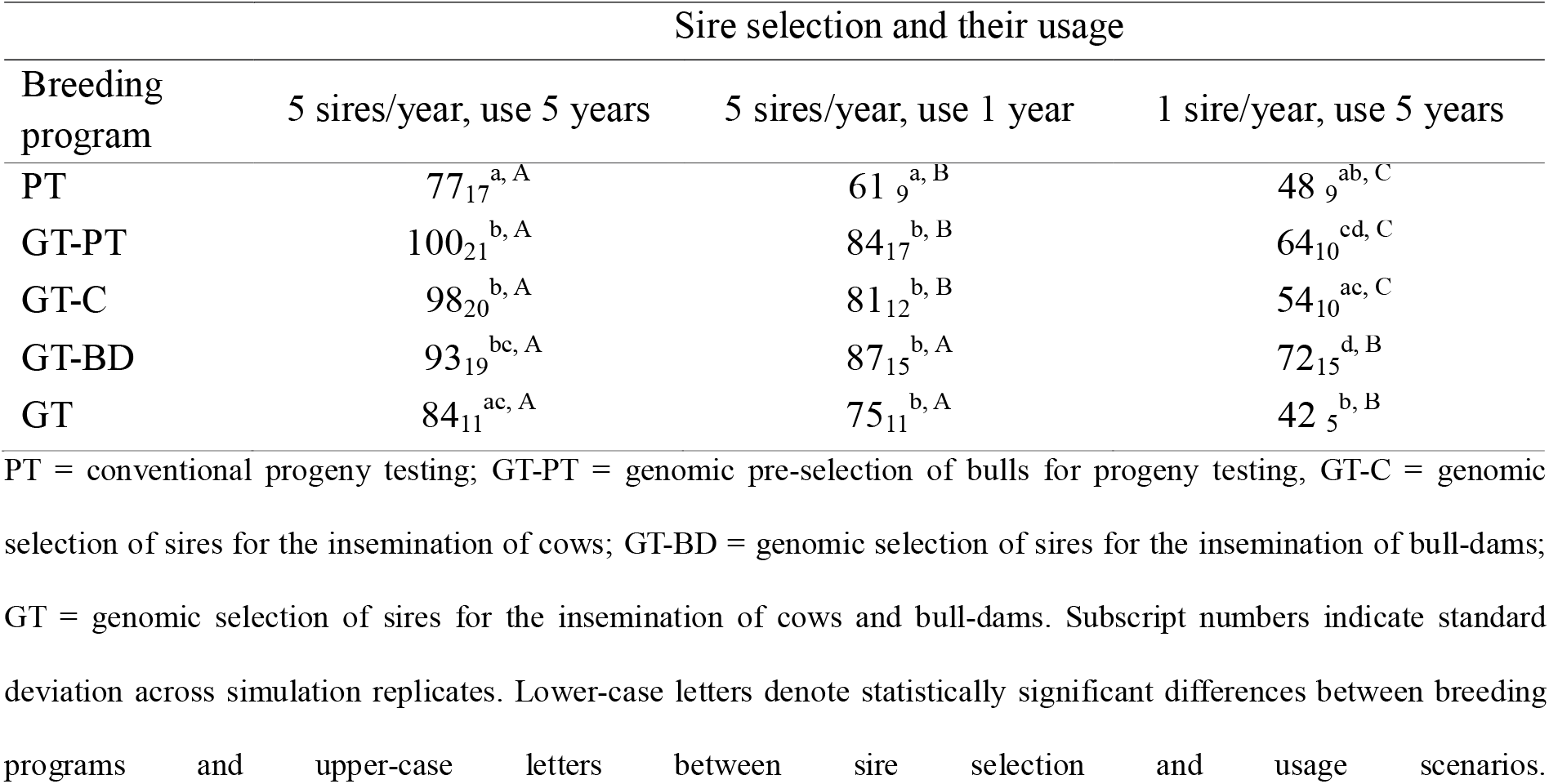
Efficiency of converting genetic variation into gain by breeding program and by sire selection and their usage scenario.

| Breeding<br>program | Sire selection and their usage |  |  |
| --- | --- | --- | --- |
|  | 5 sires/year, use 5 years | 5 sires/year, use 1 year | 1 sire/year, use 5 years |
| PT | 77<sub>17</sub><sup>a, A</sup> | 61<sub>9</sub><sup>a, B</sup> | 48<sub>9</sub><sup>ab, C</sup> |
| GT-PT | 100<sub>21</sub><sup>b, A</sup> | 84<sub>17</sub><sup>b, B</sup> | 64<sub>10</sub><sup>cd, C</sup> |
| GT-C | 98<sub>20</sub><sup>b, A</sup> | 81<sub>12</sub><sup>b, B</sup> | 54<sub>10</sub><sup>ac, C</sup> |
| GT-BD | 93<sub>19</sub><sup>bc, A</sup> | 87<sub>15</sub><sup>b, A</sup> | 72<sub>15</sub><sup>d, B</sup> |
| GT | 84<sub>11</sub><sup>ac, A</sup> | 75<sub>11</sub><sup>b, A</sup> | 42<sub>5</sub><sup>b, B</sup> |

PT = conventional progeny testing; GT-PT = genomic pre-selection of bulls for progeny testing, GT-C = genomic selection of sires for the insemination of cows; GT-BD = genomic selection of sires for the insemination of bull-dams; GT = genomic selection of sires for the insemination of cows and bull-dams. Subscript numbers indicate standard deviation across simulation replicates. Lower-case letters denote statistically significant differences between breeding programs and upper-case letters between sire selection and usage scenarios.

**Figure 2:**
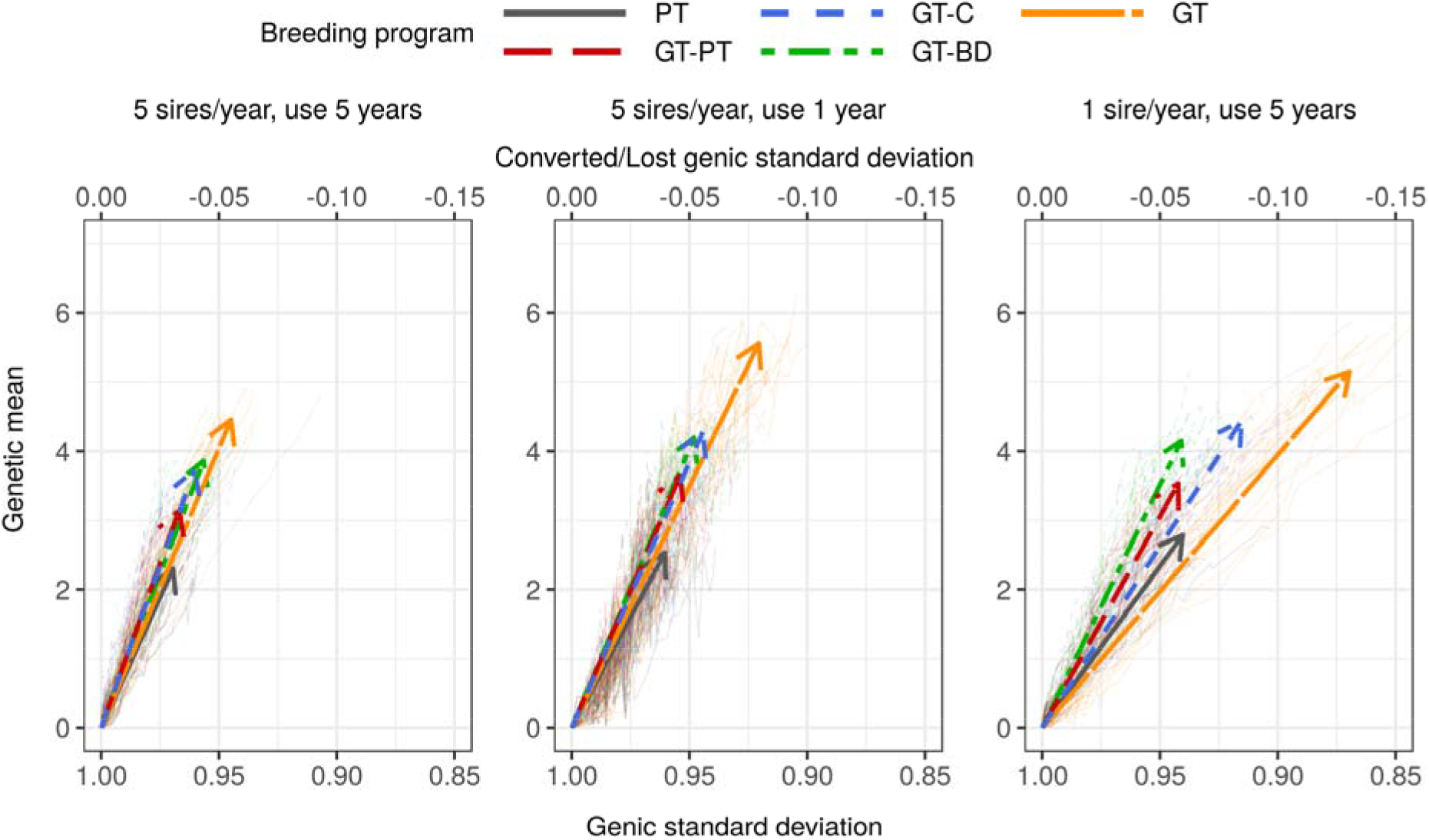
Change of genetic mean and genic standard deviation over the 20 years of selection by breeding program and by sire selection and their usage scenario. Thin lines represent individual replicates, while thick lines represent average linear regression with arrows pointing in the direction of change. PT = conventional progeny testing; GT-PT = genomic pre-selection of bulls for progeny testing, GT-C = genomic selection of sires for the insemination of cows; GT-BD = genomic selection of sires for the insemination of bull-dams; GT = genomic selection of sires for the insemination of cows and bull-dams.

### Optimum contribution selection

Optimization of male contributions increased the conversion efficiency compared to truncation selection. This is shown in Table 4 and Figure 3, which compare scenarios with truncation selection and optimum contribution selection. Optimization increased the conversion efficiency when we increased emphasis on maintenance of genetic variation. Therefore, there was always an optimum contribution selection scenario that either achieved comparable genetic gain as a truncation selection scenario, but with a smaller loss in genetic variation, or achieved larger genetic gain than a truncation selection scenario with a comparable loss in genetic variation. For example, optimum contribution selection with the target degrees of 75 achieved 21% higher genetic gain with a slightly lower rate of coancestry than the truncation selection scenario that used 5 progeny tested sires for 5 years, which taken together resulted in 121% higher conversion efficiency. Similarly, optimum contribution selection with the target degrees of 55 and 60 degrees achieved comparable or even higher genetic gain than the truncation selection scenario that used 5 genomically tested sires for 5 years on cows and bull-dams, but had slightly smaller rates of coancestry, which taken together increased conversion efficiency by respectively 38 and 51%. On the other hand, optimum contribution selection with the target degrees of 50 achieved a 26% higher genetic gain with a comparable rate of coancestry as the truncation selection scenario that used 5 genomically tested sires for 5 years. Further, optimum contribution selection with the target degrees of 45 and 50 had comparable genetic gain as the truncation selection scenario that used 5 genomically tested sires for 1 year on both, cows and bull-dams. And while the conversion efficiency for optimization at 45 degrees was comparable to the specified truncation scenario, optimization at 50 degrees had a 16% higher conversion efficiency. Increasing the emphasis on maintenance of genetic variation in optimization increased the number of selected sires and their usage over time. The average number of used sires ranged from 9.6 with the target degrees of 45 to 153.0 with the target degrees of 75. The years of usage ranged from 1.6 to 4.9 for the same span of target degrees.

**Table 4:** Comparison of breeding programs that use truncation or optimum contribution selection.

| Breeding program | Genetic gain | Sire selection accuracy | No. sires | Years in use | Generation interval (sire-sire) | Generation interval (sire-dam) | Genic standard deviation | Rate of coancestry | Effective population size | Conversion efficiency |
| --- | --- | --- | --- | --- | --- | --- | --- | --- | --- | --- |
| Truncation selection |  |  |  |  |  |  |  |  |  |  |
| 5 sires/year, use 5 years, PT | 2.50<sub>0.22</sub><sup>a</sup> | 0.88<sub>0.10</sub><sup>a</sup> | 55<sub>0.0</sub><sup>a</sup> | 2.7<sub>0.0</sub><sup>a</sup> | 9.0<sub>0.06</sub><sup>a</sup> | 7.0<sub>0.05</sub><sup>a</sup> | 0.97<sub>0.01</sub><sup>a</sup> | 0.003<sub>0.001</sub><sup>a</sup> | 172<sub>48</sub><sup>a</sup> | 77<sub>17</sub><sup>a</sup> |
| 5 sires/year, use 5 years, GT | 4.84<sub>0.26</sub><sup>b</sup> | 0.79<sub>0.12</sub><sup>b</sup> | 56<sub>0.1</sub><sup>a</sup> | 2.1<sub>0.0</sub><sup>b</sup> | 4.2<sub>0.05</sub><sup>b</sup> | 3.9<sub>0.05</sub><sup>b</sup> | 0.94<sub>0.01</sub><sup>b</sup> | 0.006<sub>0.001</sub><sup>b</sup> | 90<sub>14</sub><sup>bc</sup> | 84<sub>11</sub><sup>a</sup> |
| 5 sires/year, use 1 year, GT | 6.04<sub>0.27</sub><sup>c</sup> | 0.75<sub>0.11</sub><sup>c</sup> | 36<sub>0.0</sub><sup>b</sup> | 1.3<sub>0.0</sub><sup>c</sup> | 2.3<sub>0.05</sub><sup>c</sup> | 2.3<sub>0.00</sub><sup>c</sup> | 0.92<sub>0.01</sub><sup>cd</sup> | 0.007<sub>0.001</sub><sup>c</sup> | 72<sub>10</sub><sup>bd</sup> | 75<sub>11</sub><sup>a</sup> |
| Optimum contribution selection |  |  |  |  |  |  |  |  |  |  |
| OCS<sub>45°</sub> | 6.26<sub>0.39</sub><sup>c</sup> | 0.77<sub>0.02</sub><sup>bc</sup> | 9.6<sub>0.6</sub><sup>c</sup> | 1.6<sub>0.06</sub><sup>d</sup> | 2.8<sub>0.07</sub><sup>d</sup> | 2.7<sub>0.07</sub><sup>d</sup> | 0.91<sub>0.01</sub><sup>c</sup> | 0.008<sub>0.001</sub><sup>d</sup> | 61<sub>10</sub><sup>d</sup> | 72<sub>8</sub><sup>a</sup> |
| OCS<sub>50°</sub> | 6.10<sub>0.23</sub><sup>c</sup> | 0.79<sub>0.02</sub><sup>abc</sup> | 14.3<sub>0.9</sub><sup>c</sup> | 1.7<sub>0.05</sub><sup>d</sup> | 2.9<sub>0.07</sub><sup>e</sup> | 2.8<sub>0.05</sub><sup>e</sup> | 0.93<sub>0.01</sub><sup>d</sup> | 0.007<sub>0.001</sub><sup>bc</sup> | 75<sub>9</sub><sup>bcd</sup> | 87<sub>11</sub><sup>a</sup> |
| OCS<sub>55°</sub> | 5.27<sub>0.28</sub><sup>d</sup> | 0.79<sub>0.02</sub><sup>abc</sup> | 25.1<sub>4.7</sub><sup>d</sup> | 2.1<sub>0.17</sub><sup>b</sup> | 3.0<sub>0.07</sub><sup>f</sup> | 2.9<sub>0.06</sub><sup>f</sup> | 0.95<sub>0.01</sub><sup>be</sup> | 0.005<sub>0.001</sub><sup>e</sup> | 113<sub>22</sub><sup>ce</sup> | 115<sub>24</sub><sup>b</sup> |
| OCS<sub>60°</sub> | 4.77<sub>0.25</sub><sup>b</sup> | 0.81<sub>0.02</sub><sup>abc</sup> | 49.0<sub>8.9</sub><sup>e</sup> | 3.0<sub>0.34</sub><sup>e</sup> | 3.3<sub>0.10</sub><sup>g</sup> | 3.1<sub>0.07</sub><sup>g</sup> | 0.96<sub>0.01</sub><sup>ae</sup> | 0.004<sub>0.001</sub><sup>ae</sup> | 144<sub>24</sub><sup>e</sup> | 126<sub>17</sub><sup>b</sup> |
| OCS<sub>75°</sub> | 3.03<sub>0.17</sub><sup>e</sup> | 0.82<sub>0.01</sub><sup>abc</sup> | 153.0<sub>9.1</sub><sup>f</sup> | 4.9<sub>0.07</sub><sup>f</sup> | 4.2<sub>0.08</sub><sup>b</sup> | 4.0<sub>0.06</sub><sup>h</sup> | 0.98<sub>0.00</sub><sup>f</sup> | 0.002<sub>0.001</sub><sup>f</sup> | 276<sub>43</sub><sup>f</sup> | 162<sub>37</sub><sup>c</sup> |

PT = conventional progeny testing; GT = genomic selection of sires for the insemination of cows and bull-dams; OCS<sub>X°</sub> = optimum contribution selection of sires for the insemination of cows and bull-dams with the target degrees of X°. Subscript numbers indicate standard deviation across simulation replicates. Lower-case letters denote statistically significant differences between breeding programs.

**Figure 3:**
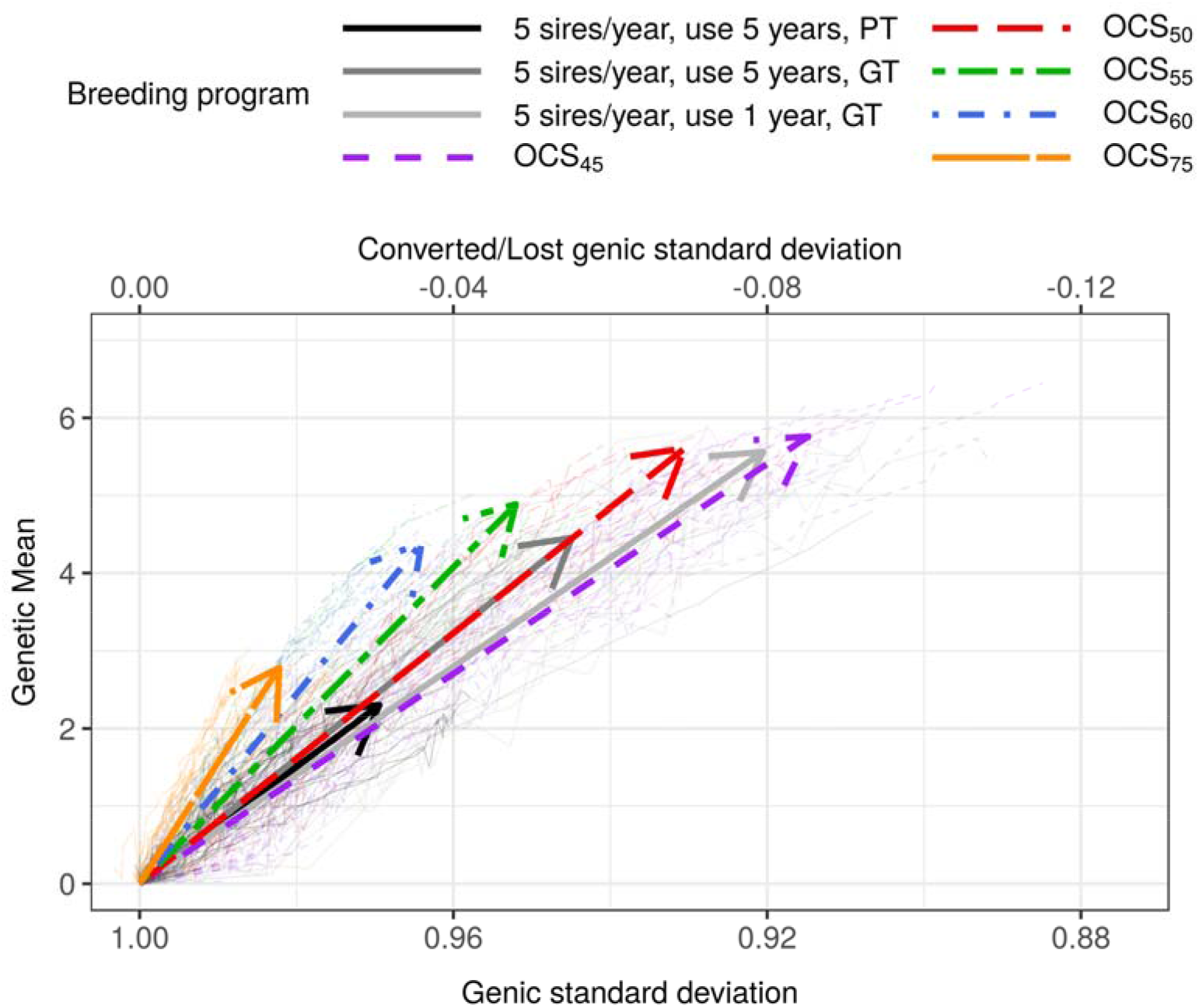
Change of genetic mean and genic standard deviation over the 20 years of selection for fixed or optimized breeding programs. Thin lines represent individual replicates, while thick lines represent average linear regression with arrows pointing in the direction of change. PT = conventional progeny testing; GT = genomic selection of sires for the insemination of cows and bull-dams; OCS_X°_ = optimum contribution selection of sires for the insemination of cows and bull-dams with the target degrees of X^°^.

## DISCUSSION

Selection dynamics in small populations differs from that of large populations. Small populations can not perform very intensive selection due to limited resources that allow for testing of only limited number of individuals. Further on, due to limited number of animals and progeny per sire, small populations struggle with the accuracy of progeny and genomic testing. And last, limited accuracy and limited number of animals could potentially affect genetic variation of the population. Despite all this, small populations have to find a way to deliver both short- and long-term genetic gain to stay competitive with larger populations and to justify domestic selection. The results show that we can increase genetic gain in such populations by implementing the genomic selection of sires, a faster turn-over of sires, and increasing the intensity of sire selection. However, these strategies also increase the loss of genetic variation, though this loss has to be assessed against the larger genetic gains. For this reason, we evaluated the efficiency of converting genetic variation into genetic gain. The results show that in small dairy populations the conversion efficiency can be improved by the simultaneous use of genomically and progeny tested sires. Optimization of male contributions can further increase the conversion efficiency. Specifically, it can increase the genetic gain of the truncation selection with a comparable loss of genetic variation or it can reduce the loss of genetic variation with a comparable genetic gain. To address these main findings, we divided discussion into four parts: i) how genomic truncation selection affects genetic gain in small populations and how this compares to large populations; ii) how genomic truncation selection affects the loss of genetic variation in small populations; iii) how optimum contribution selection can increase the conversion efficiency, which has implications for small and large populations; and iv) how small populations could further leverage the benefits of genomic selection.

### Genetic gain with genomic truncation selection

As expected, genomic selection increased the genetic gain in all sire selection and usage scenarios. This was due to a higher selection accuracy for young non-phenotyped animals and reduced generation interval (Schaeffer, 2006). Using genomic prediction as the pre-selection step increased genetic gain between 37% and 59% in different scenarios without reducing generation interval. This is a larger increase than in studies of larger populations (Pryce et al., 2010) or larger progeny groups (Lillehammer et al., 2011). In small populations additional benefit of genomic pre-selection comes from the fact that progeny testing is not as accurate as in large populations due to smaller progeny groups. Reducing the generation interval by using young genomically tested sires directly on cows and bull-dams further increased genetic gain between up to 144% when we used 5 sires per year and up to 126% when we maximized intensity and used only 1 sire per year. These results are largely in concordance with Pryce et al. (2010), Lillehammer et al. (2011) and de Roos et al. (2011), although these studies evaluated typical large cattle populations with about ten-times larger number of selection candidates.

Thomasen et al. (2014) argued that the benefit of genomic selection in small dairy populations is undermined by a limited selection accuracy for young non-phenotyped animals caused by a small reference population. A small reference population will invariably lead to inaccurate genomic predictions. In this study we achieved comparable accuracies of about 0.8 with limited progeny test and with genomic prediction based on a reference population of about 11,000 cows and 100 progeny tested sires, that was updated each year. Recent drops in prices for genome-wide genotyping should enable small dairy populations to build such reference populations. Further, some phenotyping resources could be diverted to genotyping to maximize return on investment. A comparable level of accuracy can be also achieved with international reference populations (Jorjani, 2012; Špehar et al., 2013) or a combination of national and international reference populations (Vandenplas and Gengler, 2015; Vandenplas et al., 2017; Vandenplas et al., 2018). When this level of accuracy is combined with a reduced generation interval, small populations can achieve substantially larger genetic gains than with progeny testing. Finally, increasing the selection intensity to the unrealistic use of just one sire, to come closer to the intensity of selection in large populations, further increased genetic gain, but with a considerable loss in genetic variation that started to limit genetic gain within the simulated 20 years.

### Loss of genetic variation with genomic truncation selection

The results show that small populations can increase genetic gain without increasing the loss of genetic variation by using genomic pre-selection of bulls for progeny testing. All other genomic selection scenarios increased the loss of genetic variation compared to a conventional scenario with progeny testing, although the accuracies of progeny and genomic tests were comparable and that we selected the same number of sires per year. We observed this with genic and genetic variance as well as effective population size (measured with pedigree and neutral, marker or causal loci). While losses of genic and genetic variance in the simulated period of 20 years were not substantial (at most 0.13 genic standard deviation), the changes in effective population size were substantial – from about 175 with the conventional scenarios to about 80 with the full genomic scenarios, which indicates reduced sustainability.

Our results for the rate of coancestry are not in concordance with what was observed in studies of large populations (Pryce et al., 2010) or with higher selection intensity (Lillehammer et al., 2011). which observed lower rates with genomic selection. However, lower intensity of selection in small populations stems from fewer tested animals, and not more selected, which reduces a genetic pool for selection. Our results are more in line with Doekes et al. (2018). They attribute the higher rates of inbreeding with genomic selection to the fact, that the animals with a higher relatedness to the reference population have more accurate genomic predictions and are more likely to deviate substantially and therefore to be selected (Habier et al., 2007; Clark et al., 2012). Another explanation for a larger loss of genetic variability with genomic selection is that shortening generation interval increases the turnover of germplasm from year to year, which increases genetic gain per unit of time, but also increases the loss of genetic variation per unit of time (Buch et al., 2012; Boichard et al., 2015; Gorjanc et al., 2018). Further, studies mostly report the rate of inbreeding, which measures increase in individual homozygosity (Pryce et al., 2010; Doekes et al. 2018), while we report the rate of coancestry, which measures increase in population homozygosity. While these two measures are correlated, it is the rate of coancestry that determines the sustainability of a breeding program.

To compare the simultaneous change in genetic gain and loss in genetic variation we compared different scenarios with the efficiency of converting genetic variation into genetic gain. We measured this with a linear regression of the achieved genetic gain on the lost genic standard deviation (Gorjanc et al., 2018). We found that in small cattle populations genomic pre-selection for progeny test and hybrid scenarios achieved the highest conversion efficiencies. The two extremes – conventional and complete genomic scenarios – were the least efficient. Despite their similar conversion efficiencies, there are large differences between these scenarios – namely, the genomic scenario almost doubled genetic gain. The conventional scenario had low conversion efficiency due to a small genetic gain (caused by long generation intervals) although it retained most of genetic variation. The low conversion efficiency of the conventional scenarios could be specific to small populations, since the accuracy and selection intensity of progeny testing is smaller than in large populations. The completely genomic scenario had low conversion efficiency despite a large genetic gain (caused by short generation intervals) as it lost the most of genetic variation.

Increasing the turnover of the sires and increasing selection intensity have different consequences on short and long-term success of selection. Although both of these scenarios increase genetic gain (up to 125%), increasing the intensity also increased the loss of genetic variation and in turn reduced conversion efficiency. Increased turn-over of sires from 5 to 1 year in this study achieved higher genetic gain over the 20 years than reducing the number of sires from 5 to 1, because it did not impact genetic variation so severely.

### Comparison of truncation and optimum contribution selection

Optimization of male contributions increased the conversion efficiency of truncation selection scenarios. The optimization involved all active males - the young calves with genomic prediction and sires selected in previous years – either young sires with genomic test or older with progeny test. Optimum contribution selection with genomic information has been tested before (e.g. Clark et al., 2013) with the conclusion that there is not much scope for optimization with genomic relationships unless there are very large full-sib families. Here we use optimal contribution selection to optimize selection and usage of genomically and progeny tested bulls of different ages and observe substantial differences over 20 years in a small dairy population. We achieved this by optimizing male contributions with a range of emphasis on genetic gain versus maintenance of genetic variation. In this we followed the multi-objective approach of Kinghorn (2011), where the emphasis is measured with the angle between truncation selection solution and targeted optimum contribution selection solution.

For every truncation selection scenario, we found an optimum contribution selection scenario that increased conversion efficiency. This higher efficiency was either achieved with the same genetic gain but smaller loss of genetic variation than truncation selection or with a higher genetic gain and the same loss of genetic variation as truncation selection. This improvement was achieved by optimized selection and usage of sires. For example, the average number of sires with the truncation selection of 5 progeny tested sires that were used for 5 years was about 55 (this includes young, natural service and proven bulls). Here the sires of the same age and the same status had an approximately the same number of progeny. This scenario achieved genetic gain of 2.50 genetic standard deviations, generation interval for sire-sire and sire-dam paths of 9.0 and 7.0 years, effective population size of 172 and conversion efficiency of 77. A comparable number of sires (49) was used with the optimization targeting 60 degrees, which involved mostly young sires (3 years in use). Their optimized usage delivered genetic gain of 4.77 genetic standard deviations, generation interval for sire-sire and sire-dam paths of 3.3 and 3.1 years, effective population size of 144 and conversion efficiency of 126. The highest genetic gain was achieved with the targeted degrees between 45 and 50. These targets drive optimization to achieve every year between 71% and 65% of maximum possible genetic gain with truncation selection and between 71% and 77% minimum possible group coancestry (Kinghorn, 2011; Gorjanc and Hickey, 2018). Further, although the optimization could choose genomically and progeny tested bulls, we observed that it chose mostly young genomically tested bulls, for example the maximum years in use was on average 4.9 when we optimized for 75 target degrees. This is in contrast with truncation selection scenarios, where the highest conversion efficiency was achieved with the simultaneous use of genomically and progeny tested bulls.

The results have implications also for large populations, namely they show that genomic selection is increasing turnover of germplasm per year with positive effect on genetic gain and negative effect on genetic variation. This has been already indicated in real large populations (Doekes et al., 2018). While our results are likely specific to small populations, combining these with the results from a wheat simulation study (Gorjanc et al., 2018) that used a small or a large number of parents suggest that both small and large populations can increase the conversion efficiency of genomic selection by optimizing contributions.

### Further opportunities

There are further opportunities with genomic selection for small populations that we have not addressed in this study. We specifically highlight the increasing number of genotyped females and the role of importation of external genetics. In this paper we have focused only on comparing male selection and usage strategies that required minimal changes to a breeding program. However, genotyping prices have decreased substantially in the recent years and it’s likely that in future a significant proportion of cows will be genotyped. This will increase accuracy of genetic evaluation of cows early in their lives and enable even shorter generation intervals. It will also enable accurate assessment of relationships amongst cows and bulls and open possibility for further optimization. This has been partially realized in this study by combining genomic and pedigree relationships through the single-step genetic evaluation method, which propagates all the genomic information throughout a pedigree (Legarra et al., 2009).

Many dairy breeding programs, small and large, supplement their internal breeding activities with importation of external genetics. Importation is of particular importance for small populations because they struggle to be competitive due to limited financial resources for collecting data and limited numbers of animals for collecting data and for use as selection candidates. Combining own breeding and importation opens further possibilities for optimization as it expands the genetic pool for breeding. Further, genomic selection now enables accurate genetic evaluation and relationship of foreign animals to a local population. Such foreign animals could be added into the presented optimization to exploit the expanded genetic pool and further increase sustainability of small breeding populations.

## CONCLUSION

This paper evaluated different genomic breeding programs in a small dairy cattle population with truncation selection to quantify its short- and long-term success. Furthermore, it evaluated the value of optimizing male contributions to increase efficiency of converting genetic variation into genetic gain. We concluded that genomic selection increases short-term genetic gain but can also improve long-term genetic gain when used in combination with conventional selection. We also showed that optimum contribution selection improves conversion efficiency at a comparable genetic gain or achieves higher genetic gain at a similar conversion efficiency. Our results will be of help to breeding organization that aim to implement sustainable genomic selection.

## ACKNOWLEDGMENTS

The authors acknowledge T. Perpar (Agricultural Institute of Slovenia), M. Rigler (Chamber of Agriculture and Forestry of Slovenia), and K. Potočnik (Biotechnical Faculty, University of Ljubljana) for their advice about the Brown-Swiss breeding scheme and Andres Legarra (INRA) and Ivan Pocrnic (University of Georgia) for their advice about preparing the single-step coancestry matrix. The first author also thanks the European Association for Animal Production (EAAP) for scholarship to present preliminary results of this study at the 2018 annual meeting. G. Gorjanc and J. M. Hickey acknowledge support from the BBSRC funding to The Roslin Institute (BBS/E/D/30002275).

## APPENDIX

**Table S1:** Generation interval by path of selection, by breeding program and by sire selection and their usage scenario.

| Breeding program | Sire selection and usage |  |  |  |  |  |
| --- | --- | --- | --- | --- | --- | --- |
|  | 5 sires/year, use 5 years |  | 5 sires/year, use 1 year |  | 1 sire/year, use 5 years |  |
|  | Sire of sires | Sire of dams | Sire of sires | Sire of dams | Sire of sires | Sire of dams |
| PT | 9.0<sub>0.06</sub><sup>ab,A</sup> | 7.0<sub>0.05</sub><sup>a,A</sup> | 7.1<sub>0.00</sub><sup>a,B</sup> | 5.8<sub>0.02</sub><sup>a,B</sup> | 9.1<sub>0.07</sub><sup>a,C</sup> | 7.7<sub>0.00</sub><sup>a,C</sup> |
| GT-PT | 9.0<sub>0.06</sub><sup>a,A</sup> | 7.0<sub>0.05</sub><sup>a,A</sup> | 7.1<sub>0.00</sub><sup>a,B</sup> | 5.8<sub>0.00</sub><sup>a,B</sup> | 9.1<sub>0.07</sub><sup>a,C</sup> | 7.7<sub>0.02</sub><sup>a,C</sup> |
| GT-C | 9.0<sub>0.05</sub><sup>b,A</sup> | 4.1<sub>0.04</sub><sup>b,A</sup> | 7.1<sub>0.00</sub><sup>a,B</sup> | 2.5<sub>0.00</sub><sup>b,B</sup> | 9.1<sub>0.06</sub><sup>a,C</sup> | 4.1<sub>0.00</sub><sup>b,A</sup> |
| GT-BD | 3.8<sub>0.05</sub><sup>c,A</sup> | 7.0<sub>0.05</sub><sup>c,A</sup> | 3.8<sub>0.05</sub><sup>b,A</sup> | 5.7<sub>0.00</sub><sup>c,B</sup> | 3.8<sub>0.04</sub><sup>b,A</sup> | 7.6<sub>0.00</sub><sup>c,C</sup> |
| GT | 4.2<sub>0.05</sub><sup>d,A</sup> | 3.9<sub>0.05</sub><sup>d,A</sup> | 2.3<sub>0.05</sub><sup>c,B</sup> | 2.3<sub>0.00</sub><sup>d,B</sup> | 4.2<sub>0.07</sub><sup>c,A</sup> | 3.9<sub>0.00</sub><sup>d,C</sup> |

PT = conventional progeny testing; GT-PT = genomic pre-selection of bulls for progeny testing, GT-C = genomic selection of sires for the insemination of cows; GT-BD = genomic selection of sires for the insemination of bull-dams; GT = genomic selection of sires for the insemination of cows and bull-dams. Subscript numbers indicate standard deviation across simulation replicates. Lower-case letters denote statistically significant differences between breeding programs and upper-case letters between sire selection and usage scenarios.

**Table S2:** Accuracy of selection and prediction bias by animal category, by breeding program and by sire selection and their usage scenario.

| Breeding<br>program | Sire selection and usage |  |  |  |  |  |
| --- | --- | --- | --- | --- | --- | --- |
|  | 5 sires/year, use 5 years |  | 5 sires/year, use 1 year |  | 1 sire/year, use 5 years |  |
|  | Accuracy | Bias | Accuracy | Bias | Accuracy | Bias |
| PT |  |  |  |  |  |  |
| young bulls | 0.31<sub>0.18</sub> | 0.17<sub>0.11</sub> | 0.23<sub>0.20</sub> | 0.12<sub>0.11</sub> | 0.30<sub>0.19</sub> | 0.17<sub>0.12</sub> |
| sires | 0.88<sub>0.10</sub> | 0.83<sub>0.18</sub> | 0.87<sub>0.10</sub> | 0.82<sub>0.18</sub> | 0.82<sub>0.17</sub> | 0.76<sub>0.22</sub> |
| heifers | 0.35<sub>0.05</sub> | 0.22<sub>0.03</sub> | 0.32<sub>0.06</sub> | 0.19<sub>0.05</sub> | 0.36<sub>0.06</sub> | 0.22<sub>0.05</sub> |
| GT-PT |  |  |  |  |  |  |
| young bulls | 0.80<sub>0.11</sub> | 1.01<sub>0.22</sub> | 0.78<sub>0.12</sub> | 1.01<sub>0.23</sub> | 0.80<sub>0.11</sub> | 1.06<sub>0.23</sub> |
| sires | 0.82<sub>0.16</sub> | 0.92<sub>0.27</sub> | 0.83<sub>0.15</sub> | 0.92<sub>0.25</sub> | 0.78<sub>0.20</sub> | 0.88<sub>0.31</sub> |
| heifers | 0.47<sub>0.04</sub> | 0.34<sub>0.05</sub> | 0.44<sub>0.05</sub> | 0.30<sub>0.05</sub> | 0.46<sub>0.06</sub> | 0.33<sub>0.06</sub> |
| GT-C |  |  |  |  |  |  |
| young bulls | 0.81<sub>0.11</sub> | 1.02<sub>0.21</sub> | 0.80<sub>0.11</sub> | 1.03<sub>0.22</sub> | 0.82<sub>0.12</sub> | 1.07<sub>0.23</sub> |
| sires | 0.89<sub>0.09</sub> | 0.93<sub>0.18</sub> | 0.90<sub>0.08</sub> | 0.94<sub>0.21</sub> | 0.87<sub>0.13</sub> | 0.89<sub>0.20</sub> |
| heifers | 0.44<sub>0.05</sub> | 0.31<sub>0.05</sub> | 0.41<sub>0.05</sub> | 0.27<sub>0.05</sub> | 0.46<sub>0.07</sub> | 0.34<sub>0.08</sub> |
| GT-BD |  |  |  |  |  |  |
| young bulls | 0.77<sub>0.11</sub> | 1.00<sub>0.22</sub> | 0.78<sub>0.12</sub> | 1.00<sub>0.23</sub> | 0.77<sub>0.11</sub> | 0.98<sub>0.22</sub> |
| sires | 0.84<sub>0.14</sub> | 0.92<sub>0.23</sub> | 0.82<sub>0.16</sub> | 0.92<sub>0.29</sub> | 0.80<sub>0.17</sub> | 0.89<sub>0.27</sub> |
| heifers | 0.51<sub>0.06</sub> | 0.39<sub>0.07</sub> | 0.46<sub>0.06</sub> | 0.32<sub>0.06</sub> | 0.49<sub>0.06</sub> | 0.36<sub>0.06</sub> |
| GT |  |  |  |  |  |  |
| young bulls | 0.79<sub>0.12</sub> | 1.03<sub>0.22</sub> | 0.75<sub>0.11</sub> | 0.96<sub>0.22</sub> | 0.80<sub>0.12</sub> | 1.16<sub>0.30</sub> |
| heifers | 0.48<sub>0.05</sub> | 0.36<sub>0.06</sub> | 0.43<sub>0.06</sub> | 0.30<sub>0.05</sub> | 0.49<sub>0.07</sub> | 0.39<sub>0.13</sub> |

PT = conventional progeny testing; GT-PT = genomic pre-selection of bulls for progeny testing, GT-C = genomic selection of sires for the insemination of cows; GT-BD = genomic selection of sires for the insemination of bull-dams; GT = genomic selection of sires for the insemination of cows and bull-dams. Subscript numbers indicate standard deviation across simulation replicates.

**Figure S1:**
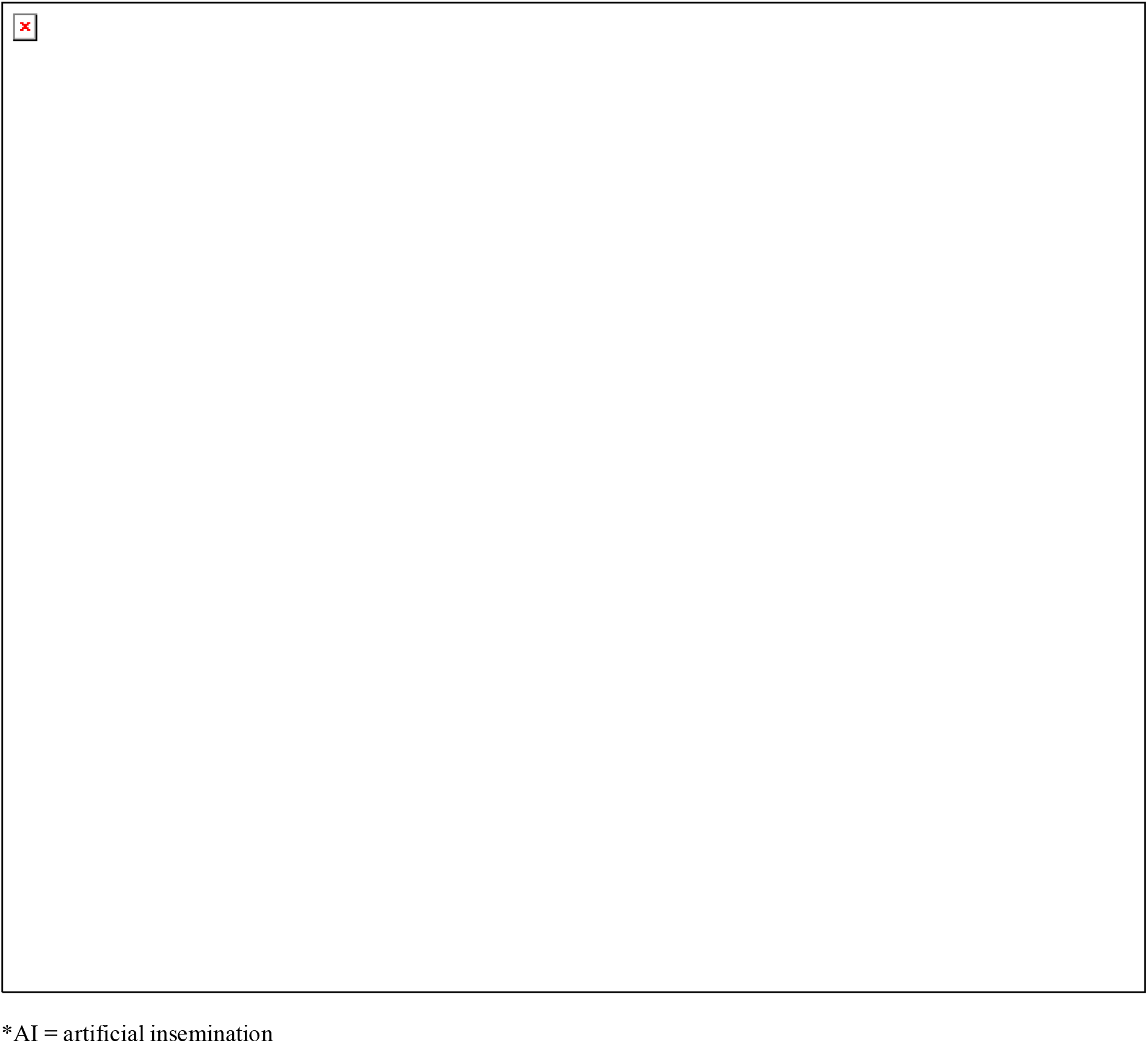
Description of the population structure and selection procedure of the simulated population for females. The arrows represent selection decisions and the numbers in bold represent the number of animals in each category.

**Figure S2:**
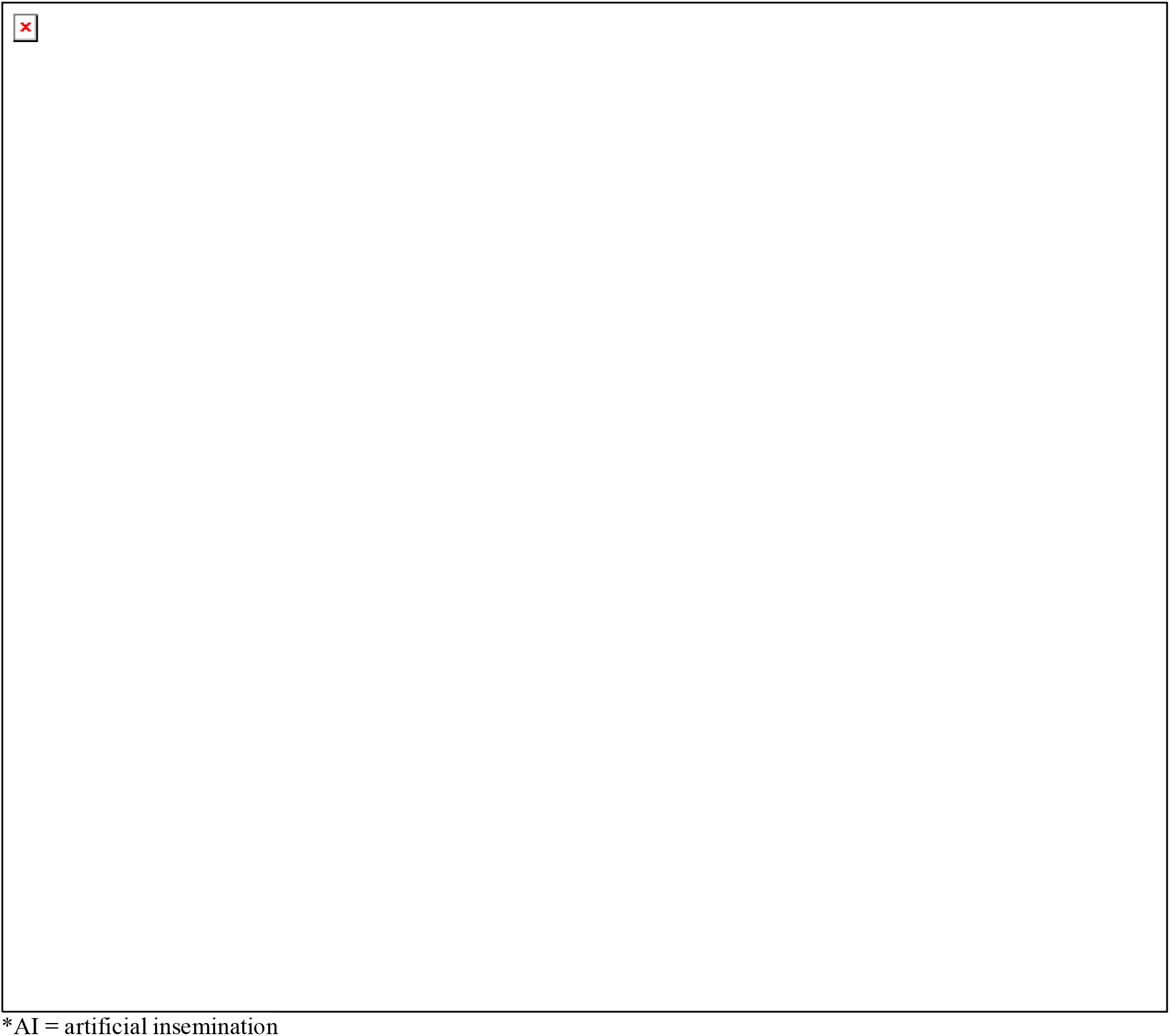
Description of the population structure and selection procedure of the simulated population for progeny tested males. The arrows represent selection decisions and the numbers in bold represent the number of animals in each category.

**Figure S3:**
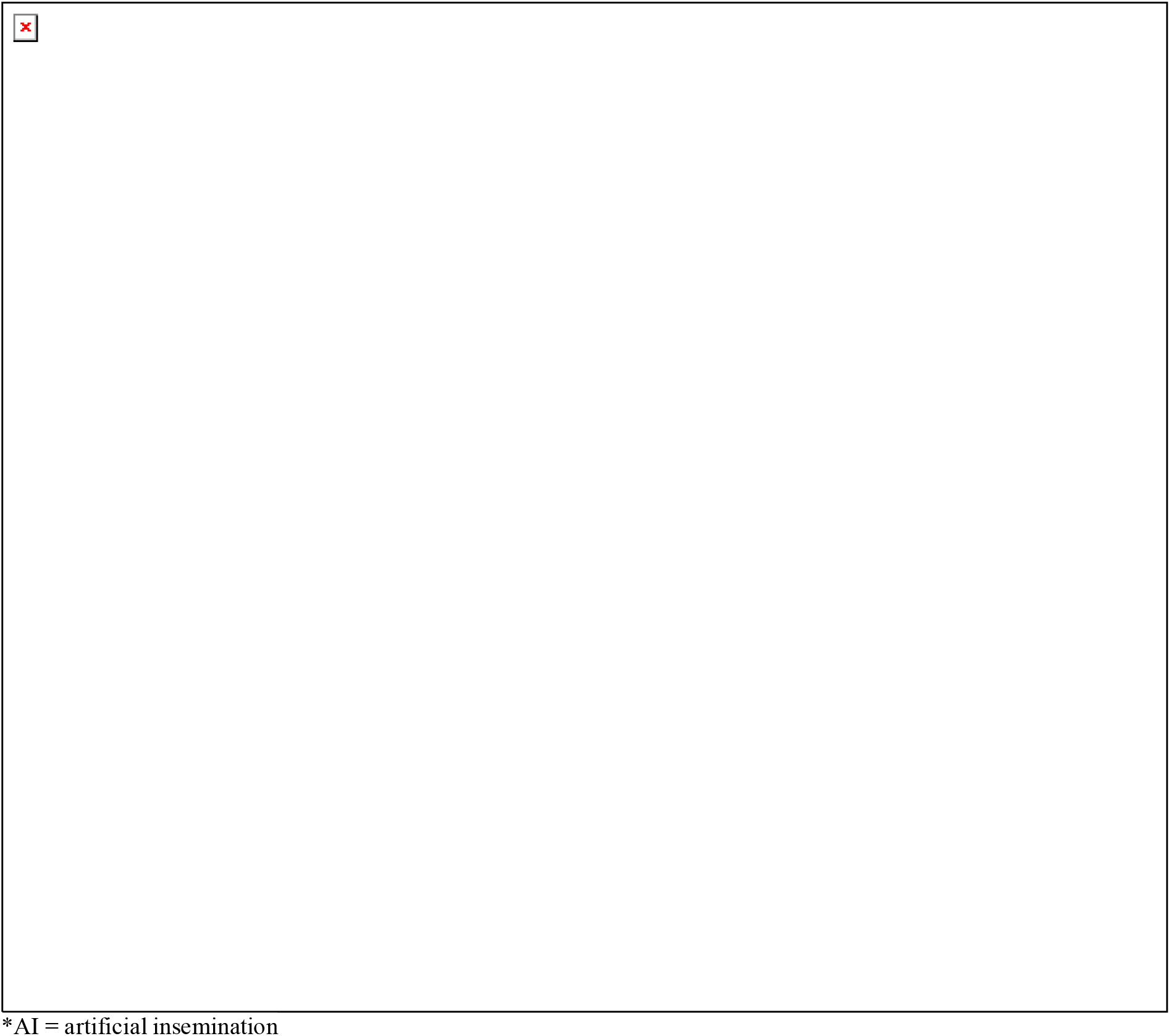
Description of the population structure and selection procedure of the simulated population for genomically tested males. The arrows represent selection decisions and the numbers in bold represent the number of animals in each category.

